# Sharp wave ripple coupling in zebrafish hippocampus and basolateral amygdala

**DOI:** 10.1101/2023.02.07.527487

**Authors:** I. Blanco, A. Caccavano, J. Wu, S. Vicini, E. Glasgow, K. Conant

## Abstract

The mammalian hippocampus exhibits sharp wave events (1-30 Hz) with an often-present superimposed fast ripple oscillation (120-200 Hz) forming a sharp wave ripple (SWR) complex. During slow wave sleep or consummatory behaviors, SWRs result from the sequential spiking of hippocampal cell assemblies initially activated during imagined or learned experiences. SWRs occur in tandem with cortical/subcortical assemblies critical to the long-term storage of specific memory types. Leveraging juvenile zebrafish, we show that SWR events in their hippocampal homologue, the anterodorsolateral lobe (ADL), in *ex vivo* whole-brains are locally generated and maintained. SWR events were also recorded in the basolateral amygdala (BLA). Concomitant single cell calcium imaging and local field potential (LFP) recordings showed that BLA SWs couple to ADL SWs. Calcium imaging recordings of whole-brains demonstrated that ADL and BLA SWRs are endogenously and spontaneously silenced by the activation of a more caudal population of putative cholinergic cells. Electrical stimulation of this caudal region silenced ADL SWs. Our results suggest that the SWR-generating circuit is evolutionarily conserved through shared acetylcholine modulating mechanisms. These findings further our understanding of neuronal population dynamics in the zebrafish brain and highlights their advantage for simultaneously recording SW/SWRs and single cell activity in diverse brain regions.

## INTRODUCTION

The mammalian hippocampus exhibits intrinsic spontaneous neuronal population oscillations known as sharp wave ripples (SWRs) during periods of awake immobility and slow wave sleep (Buzsáki et al., 1983; Buzsáki, 1986). SWRs are characterized by a relatively slow (1-30 Hz) strong deflection or sharp wave (SW), with an often-present superimposed fast ripple oscillation (150-250 Hz for rodents and 80-140 Hz for humans). SWRs are associated with the rapid and sequential replay of hippocampal place cell assemblies previously activated during learning experiences (Joo & Frank, 2018) and are necessary for the transfer of experiences to the neocortex for memory consolidation (Axmacher et al., 2006; Buzsaki et al., 1992; Buzsáki, 1986, 1996, 2015; Colgin, 2016; Ego-Stengel & Wilson, 2009; Fell et al., 2001; Girardeau et al., 2009, 2014; Jones et al., 2019; Mölle et al., 2006; Sadowski et al., 2016; Schlingloff et al., 2014; Skelin et al., 2021). Electrical or optogenetic disruption of SWRs is associated with impaired memory consolidation and recall (Ego-Stengel & Wilson, 2009; Girardeau et al., 2009). The basolateral amygdala (BLA), a subcortical brain region crucial for the processing, consolidation, and recall of emotional memories (McGaugh, 2004; O’Neill et al., 2018; Pignatelli & Beyeler, 2019), specifically associative and fear memories (Bocchio et al., 2017; Stork & Pape, 2002), also exhibits oscillatory neuronal patterns, including SWRs in mammals (Paré, 2002; Perumal et al., 2021; Ponomarenko et al., 2003; Popescu & Paré, 2011). Human studies have corroborated the translational relevance of using mammal animal models for recording and measuring SWRs events by showing mammalian SWR events’ importance and occurrence during memory consolidation and recall (Feng et al., 2018; Norman et al., 2019; Vaz et al., 2020). Given the role of the hippocampus and amygdala in learning and memory consolidation, it can be postulated that the consolidation of hippocampal-dependent memories accompanied by emotional valence requires the occurrence of time-locked SWR events between both areas. In fact, Cox *et al*., 2020 showed that in humans, amygdala SWRs are temporally coupled with hippocampal SWRs during non-rapid eye movement (NREM) sleep.

SWRs are modulated by neurotransmitters including GABA, glutamate, and acetylcholine (Buzsáki et al., 1983; Buzsáki, 1996, 2015; Schlingloff et al., 2014; Sullivan et al., 2011; Sun et al., 2018; Ylinen et al., 1995; Y. Zhang et al., 2021). SWR events are the result of the interplay between excitation and inhibition of neuronal subpopulations in the hippocampus (Buzsáki, 2015; Eller et al., 2015; Schlingloff et al., 2014; Sipilä et al., 2006; Ylinen et al., 1995). The same hippocampal circuitry that gives rise to SWR events during slow wave sleep and consummatory behaviors is necessary for the occurrence of theta/gamma burst (TGB) activity during exploration and memory encoding. Transition between the two states is modulated by acetylcholine levels. Exploration and memory encoding are associated with a high cholinergic tone and the sequential firing of place cells gives rise to TGB activity. TGB activity later transitions to SWR events, during which sequential re-activation of place cell assemblies occurs in an environment of low cholinergic tone (Buzsaki et al., 1992; Buzsáki, 1996; Csicsvari & Dupret, 2014; Hasselmo, 1999; Herweg et al., 2020; Ma et al., 2020; Nyhus & Curran, 2010; O’Keefe, 1993; Osipova et al., 2006; Sederberg et al., 2006; Sullivan et al., 2011; Wilson & McNaughton, 1994; Y. Zhang et al., 2021). Studies with isolated murine hippocampal slices, which lack septal cholinergic input, have shown that increasing acetylcholine tone, exogenously, is accompanied by a decrease in the abundance of SWR events. Bath application of the muscarinic and nicotinic agonist, carbachol, abolishes SWR complexes and allows the transition to gamma and theta oscillations (Fisahn et al., 1998; Fischer et al., 2014, p.; Konopacki et al., 1987; Li et al., 2019; S. Zhang et al., 2000). Alternatively, the cholinergic antagonist, atropine, reverses these changes and increases SWR abundance in mice (Fischer et al., 2014; Hashimoto et al., 2017). Furthermore, excessive cholinergic inhibition of hippocampal SWRs can impair memory (Buzsáki, 2015; Y. Zhang et al., 2021).

SWRs have been described in the anterodorsolateral lobe (ADL) – hippocampus homologue – in *ex vivo* whole-brain preparations from adult zebrafish (Vargas et al., 2011, 2012). However, the neural substrates of juvenile zebrafish SWRs, including their spatial generation and maintenance, remain largely unknown. The homologue of the BLA can be found in the dorsal telencephalon of the zebrafish medially to the ADL. (Bartoszek et al., 2021; Ganz et al., 2015; Lal & Kawakami, 2022; Porter & Mueller, 2020; von Trotha et al., 2014). To date, it is still unknown whether the BLA of zebrafish exhibit SWRs and if the coupling between hippocampal and BLA SWRs is conserved to aid in hippocampal-dependent emotional memory consolidation in this animal model. It is also unclear whether the cholinergic system plays a role in modulating zebrafish SWR events as it does in mammals.

In this study we recorded local field potentials (LFP) from both whole-brain and de-tectomized (telencephalon-only) *ex vivo* preparations from juvenile zebrafish to determine SWR electrophysiological properties, including their modulation by different neurotransmitters. We also took advantage of the everted zebrafish brain, in which both the ADL and BLA are in the most dorsal side of the telencephalon and used concomitant LFP and calcium imaging in whole-brain *ex vivo* preparations to look at SW coupling between these regions. We found that the zebrafish SWR-like activity is locally generated and maintained in the telencephalon though its abundance and duration seem to be further controlled by extra-telencephalic afferents. Like SWRs from murine hippocampal slices and adult zebrafish, juvenile zebrafish SWs are sensitive to AMPA and GABA_A_ but not NMDA blockade (Behrens et al., 2005; Maier et al., 2003; Schlingloff et al., 2014; Vargas et al., 2012; Wu et al., 2005). Consistent with previous literature, fluctuations in cholinergic tone modulated the abundance of SWs (Vandecasteele et al., 2014; Y. Zhang et al., 2021; Zhou et al., 2019). The endogenous activation of a caudal region with putative cholinergic neurons spontaneously and transiently suppressed SWs and the electrical stimulation of this region corroborated the silencing of these events. Our results suggest that the zebrafish hippocampal homologue can locally generate and maintain SWs and that these are in turn modulated by the cholinergic system. The data also show the coupling between ADL and BLA SW events. SWR coupling between these regions supports zebrafish as a model for studying emotional memory encoding and consolidation.

## METHODS

### Zebrafish Maintenance and Husbandry

Groups of 10-20 wild-type and *Tg(elevI3:Hsa:H2B:GCaMP6s)jf5* juvenile (30-57 days old) zebrafish were housed in 2L tanks with water temperature kept at 28 °C. Males and females were not separated for this study: they were treated as one homogenous population. Zebrafish were fed twice a day and kept on a 14/10 light-dark cycle (lights on 9 AM and lights off at 11 PM). All procedures were performed in accordance with the Institutional Animal Care and Use Committee of Georgetown University, Washington DC, USA.

### Zebrafish Local Field Potential Recordings and Ca^2+^ Imaging

Zebrafish were submerged in an overdose of tricaine methanesulfonate (MS222, Sigma-Aldrich St. Louis, MO, USA) and whole-brains with the olfactory bulb intact, after decapitation and carefully removing the skull, were extracted. Brains were incubated in a chamber containing oxygenated (95% O_2_, 5% CO_2_) artificial cerebrospinal fluid (aCSF) previously described (Brenet et al., 2019), containing in mM: 134 NaCl, 2.9 KCl, 2.1 CaCl_2_, 1.2 MgCl_2_, 10 HEPES, and 10 Glucose, pH 7.4 for at least 30 minutes at room temperature throughout the recovery period and recordings: 20 to 90 minutes. Recordings were obtained from the anterodorsolateral (ADL) lobe of the telencephalon of zebrafish, previously described (Vargas et al., 2012) to exhibit spontaneous events characterized as sharp wave (SW) with ripple embedded (SWR) similar to mammals. SWRs were also recorded from the BLA. For LFP recordings, electrodes from borosilicate glass pipettes were pulled using a Sutter P87 puller with 5 controlled pulls, resulting in an approximate 150 KΩ tip resistance and were filled with aCSF. The brains were constantly perfused inside a submerged recording chamber at a high flow rate (20 mL/min) at room temperature.

For concomitant LFP recordings and Ca^2+^ imaging, *ex vivo* whole-brains from juvenile *Tg(elevI3:Hsa:H2B:GCaMP6s)jf5* zebrafish were kept in oxygenated aCSF at room temperature to recover for a minimum of 30 minutes. Brains were moved to an upright laser scanning confocal microscope system (Thor Imaging Systems Division) with constant perfusion and held down using a SHD-42/15 WI 64-1420 Harp (Warner Instruments) for recording both LFP and single cell Ca^2+^ transient activity in the telencephalon. We used the 488 nm laser (green) for Ca^2+^ transient activity recordings and videos were captured using 512×512 pixel frames at a sample rate of 10 Hz. We used a 20x objective and an imaging field of 100-150 mm to ensure complete view of the telencephalon while retaining single cell resolution.

### Local Field Potential and Calcium Imaging Analysis

Analysis of LFP recordings was performed using a custom MATLAB script previously described (Caccavano et al., 2020), with some modifications. After correcting for 60 Hz line noise and harmonics, a gaussian finite impulse response band-pass filter with corrected phase delay was applied between 1-1000 Hz and subsequently between the frequencies of interest including the SW (1-30 Hz) and ripple (120-220 Hz). The root mean square (RMS) of the SW and ripple was computed in sliding 30 and 5 ms windows, respectively. SW and ripple events were detected from the RMS signals that exceeded 6 and 4 standard deviations (SD) above baseline, while the event start and end times were determined when the RMS signals exceeded 4 and 3 SD above baseline, respectively. SWR events were defined as a subset of SW events with a concurrent ripple. For both SWs and ripples, events with a duration of less than 25 ms were discarded, and successive events with less than 150 ms inter-event interval were treated as one continuous event. To determine the baseline SD for each RMS signal in a way that accounts for both inactive and active recordings, the entire time series was binned into a histogram and a two-term gaussian fitting technique was employed to estimate both the baseline noise and true signal.

Calcium imaging data were first preprocessed with custom ImageJ (FIJI) macros previously described (Caccavano et al., 2020), before analysis in MATLAB. Raw image stacks were corrected for photobleaching, which was modeled as an exponential decay. Circular region-of-interests (ROIs) were manually placed around cell somas. For regional ROIs, neuronal clusters were chosen based on the pattern of activation before and during a silent period. The change in fluorescence over total fluorescence (Δ*F/F*) was then computed as (*F-F_0_*)/(*F_0_-Fb*), where *F_0_* was defined as the average of the ten lowest intensities for each ROI, and *F_b_* was defined as the lowest intensity pixel across the entire image and time series. These ROI time series were subsequently imported into MATLAB for coincidence detection with the LFP. Slow changes in the baseline fluorescence were corrected with a smoothed moving average spanning 25% the duration of the file. Δ*F/F* signals were then interpolated to a 2 ms sampling rate, while the LFP was down-sampled to the same common sampling rate. Calcium transients were detected above 6 SD of each Δ*F/F* signal, with event start and end times determined when the Δ*F/F* exceeded 4 SD. The baseline SD and event detection was carried out by the same algorithm as described for the SW and ripple detection.

### Pharmacological Agents

Juvenile zebrafish *ex vivo* whole-brain preparations were exposed to pharmacological agents previously shown to modulate SW events in both mammals and adult zebrafish. AP-5 was used to block N-methyl-D-aspartate (NMDA) transmission and NBQX was used to block α-amino-3-hydroxy-5-methylisoxazole-4-propionic acid (AMPA) transmission. Bicuculine Methiodide (BMI) was used to block GABA_A_ receptors. Atropine, a muscarinic receptor antagonist, and Mecamylamine (MEC), a non-competitive nicotinic receptor antagonist, were used to block cholinergic transmission. All drugs were purchased from Sigma, St. Louis, MO, USA, and stock concentrations were dissolved in distilled water. LFP activity was recorded after final bath concentrations of 30 μM AP-5, 5 μM NBQX, 5 μM NBQX + 25 μM AP-5, 30 μM Bicuculine, 10 μM atropine, and 20 μM MEC.

### Electrical Stimulation

Region 4 (PMPa) was electrically stimulated (n = 3 whole-brain preparations) with a glass microelectrode with a tip opening of 10-50 μm (~50 KΩ). The stimulus pulse duration was 0.05 ms (generated by a Master 8 stimulator), 100 pulses at 50 Hz were used. The stimulus intensity was 1-5 μA.

### Statistical Analysis

All statistical analyses were performed in GraphPad Prism 9.3. Data were tested for normality and lognormality via Shapiro-Wilk tests. Parametric Paired (Figure 2 & 5) and Unpaired Student’s t tests (Figures 1) were performed. Parametric one-way ANOVA tests (Figure 7) with post-hoc Tukey’s multiple comparisons corrections was performed after normality was accessed. Kolmogorov-Smirnov D test (Figure 2) and Chi-Square (Figure 6) were also performed.

**Figure 1.**
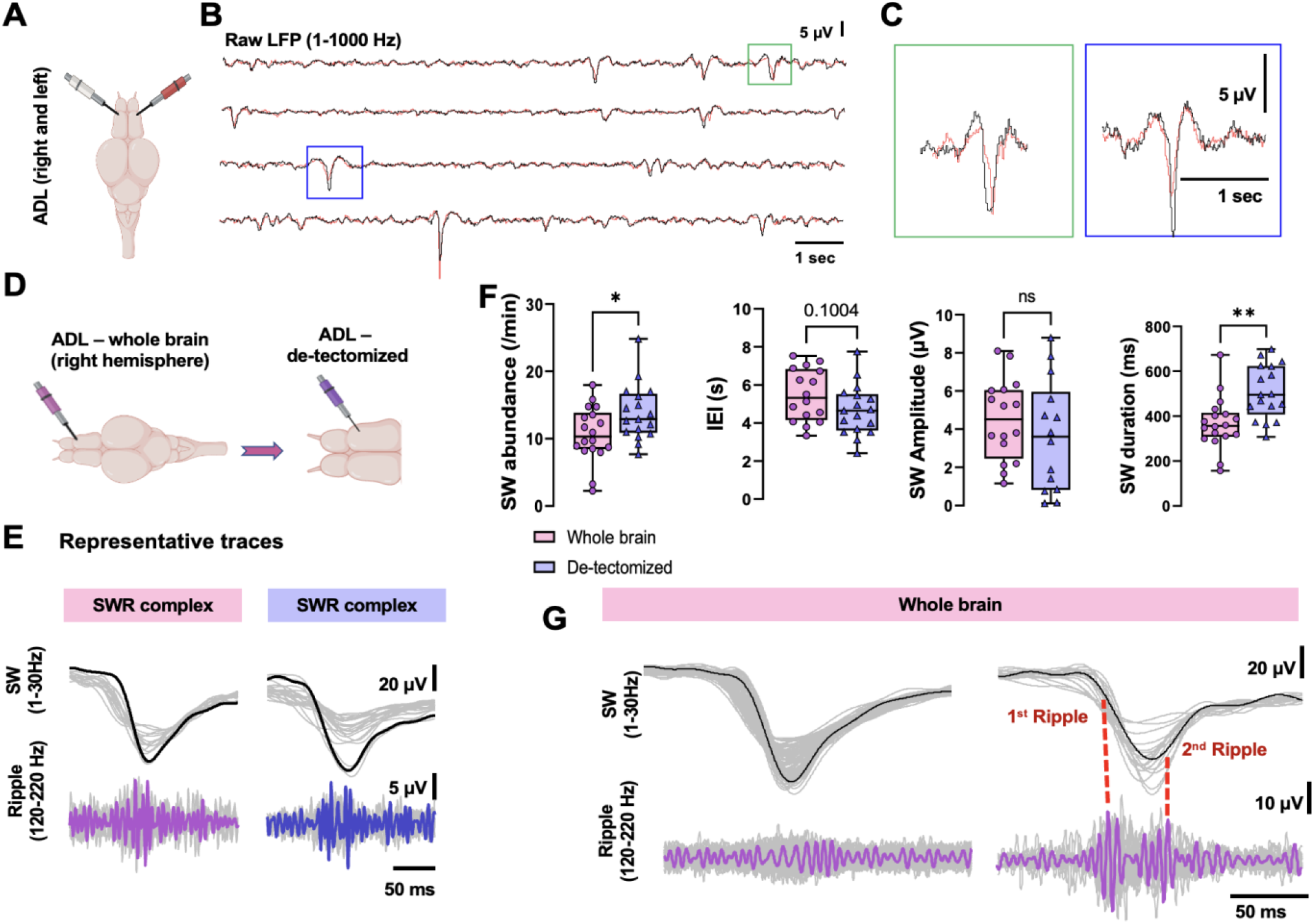
SWR events are locally generated and maintained within the telencephalon of juvenile zebrafish. (a) Representative cartoon of the zebrafish brain and electrode placement in either the right or left ADL. (b) Representative superimposed LFP traces (1-1000 Hz) from the right ADL (white) and left ADL (red). (c) Zoomed in of individual LFP oscillation from (b). (d) Representative traces of electrode placement in the ADL of whole-brain preparations (left) and de-tectomized preparations (right). (e) Representative SWR complex from whole-brains (pink) and de-dectomized preparations (dark blue). (f) Quantification of SW events, IEI, amplitude, and duration between whole-brain and de-tectomized recordings. (g) Left panel: Representative SW event with no superimposed ripple. Right panel: Representative SW event with a ripple doublet. Data analyzed by Unpaired Student’s t test. Box & whiskers plot. *p < 0.05, **p < 0.005.

**Figure 2.**
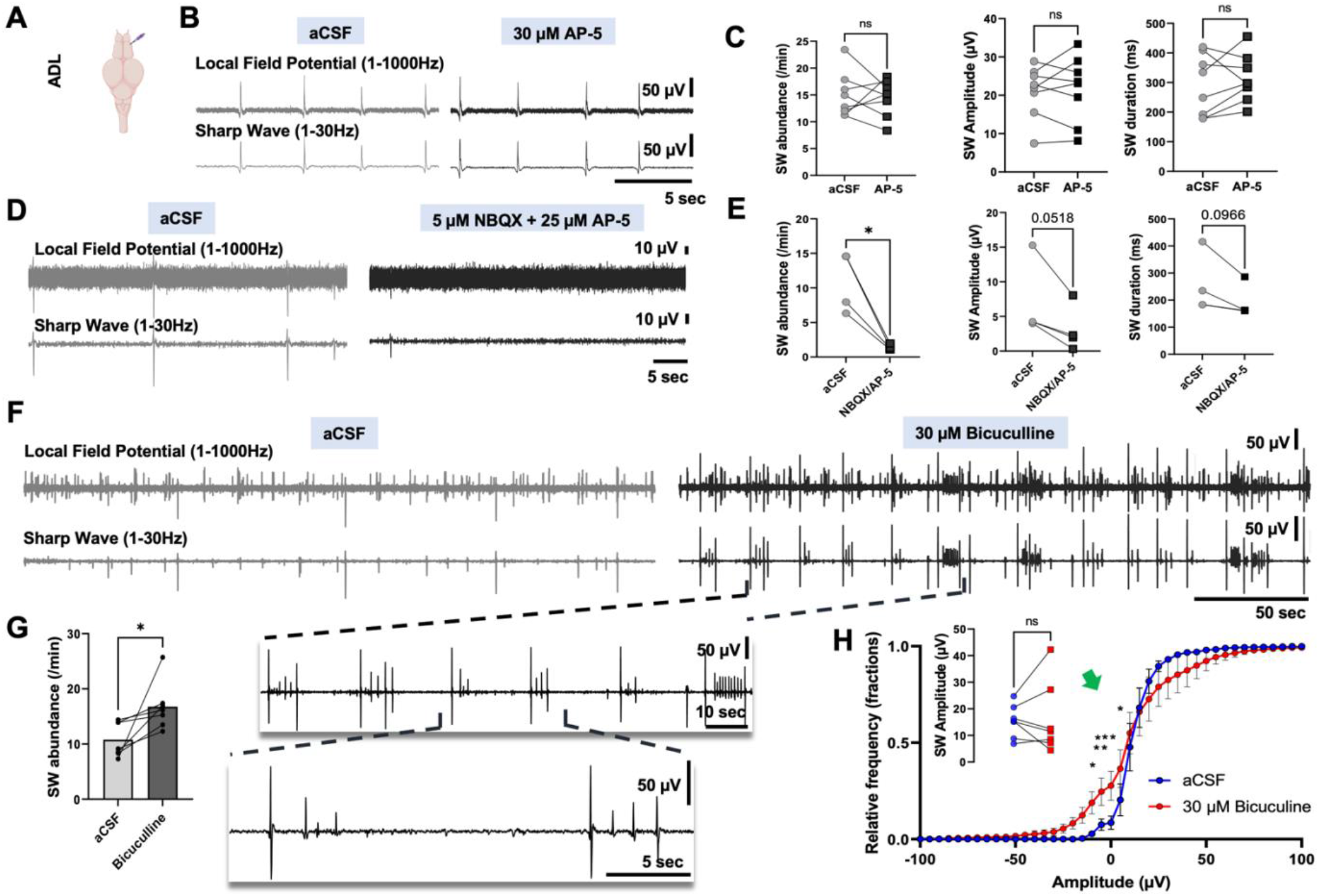
SW events in the ADL are modulated by GABA_A_ and AMPA but not by NMDA signaling. (a) Representative image of electrode placement for LFP recording from the ADL. (b-c) Exposure to 30 μM AP-5 did not change the abundance of SW events or duration. (d-e) 5 μM NBQX + 25 μM AP-5 significantly decrease the abundance of SW events and trended towards a decrease in SW event amplitude and duration. (f-g) 30 μM Bicuculline significantly increased the average abundance of SW events. (h – inset) Average amplitude after bicuculine was not changed. A significant change in the cumulative frequency was observed with a 2way ANOVA showing a shift to the left in the amplitude of SWs in the negative deflection (green arrow). *p < 0.05, **p < 0.005, ***p < 0.0005

**Figure 3.**
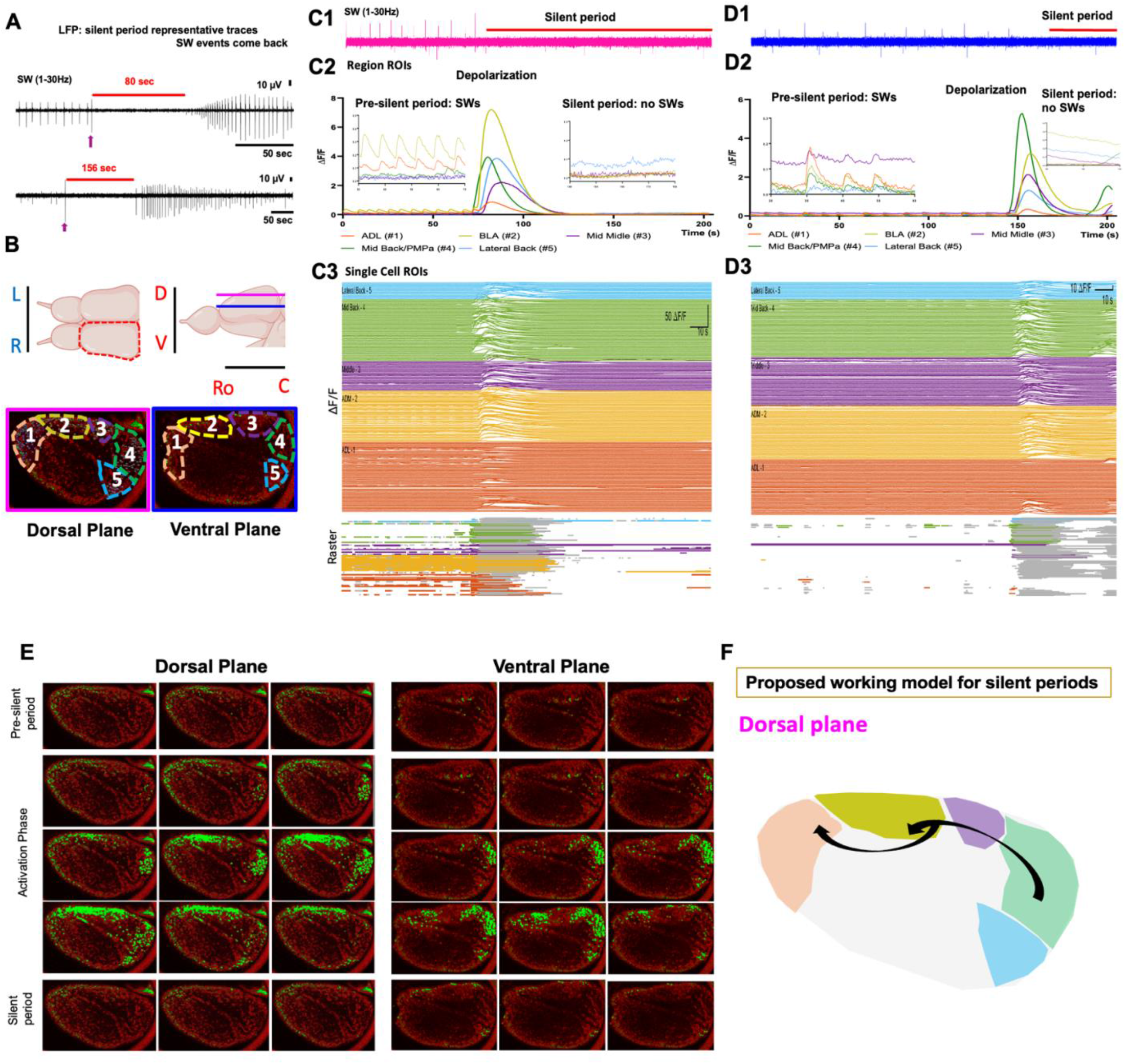
Dual LFP and calcium transient recordings reveal silent period events in the zebrafish brain following massive caudal-rostral neuronal activation. (a) Representative SW traces from juvenile zebrafish with silent events lasting from 80 to 156 seconds. Returning SW events have distinct amplitude patterns: increase, decrease, and no change. Purple arrow shows a high amplitude event that often happens before a silent period. (b) *Top:* A dorsal view drawing of a zebrafish brain showing the olfactory bulb and the telencephalon with the right telencephalon bordered by a red dashed line, and a lateral view drawing showing the olfactory bulb and the telencephalon with the approximate levels of the dorsal (pink) and ventral (blue) planes indicated with lines. Directional axes are indicated as left (L) vs right (R,), dorsal (D) vs ventral (V), and rostral (Ro) vs caudal (C). *Bottom:* Images of the left telencephalon of *Tg(elevI3:Hsa.H2B:GCaMP6s*) juvenile zebrafish under the confocal microscope with the pink bordered image corresponding to the dorsal plane and the blue bordered image to the ventral plane (about 30-50 microns apart), the medial side is facing up and the lateral side is facing down. Images are divided into five different regions of interest based on the neuronal firing associated with both the generation of SW and the massive activation preceding silent periods. (c1-d3) The same analysis and similar neuronal firing were done for the dorsal area of the telencephalon (pink) and the ventral (blue) area with regards to silent period generation. Dual recordings of neuronal population recordings (c1, d1) and regional/single cell calcium imaging (c2, d2, c3, d3) in the *Tg(elevI3:Hsa.H2B:GCaMP6s*) were aligned showing temporally coordinated firing between the ADL (region 1) and BLA (region 2) during SW events while regions 4 and 5 were silent. (c2, d2) Massive increase in the ΔF/F in all regions starting in region 4 in the dorsal and ventral regions of the telencephalon. (c2, d2 insets) Zoomed in graphs before and after the silent period. (e) Example timelapse images of the right hemisphere from the dorsal and ventral planes of a *Tg(elevI3:Hsa.H2B:GCaMP6s*) transgenic brain exhibiting SW events and a silent period – calcium transients are shown in green. During the *Pre-silent period:* regions 1 and 2 show SW events and changes in ΔF/F. During the *Activation* phase, there are no detectable SW events but a massive activation that starts in region 4 and propagates rostrally. No single cell activity during the *Silent period* phase. (f) A proposed model of caudal-to-rostral inhibition of SW events (n = 3).

**Figure 4.**
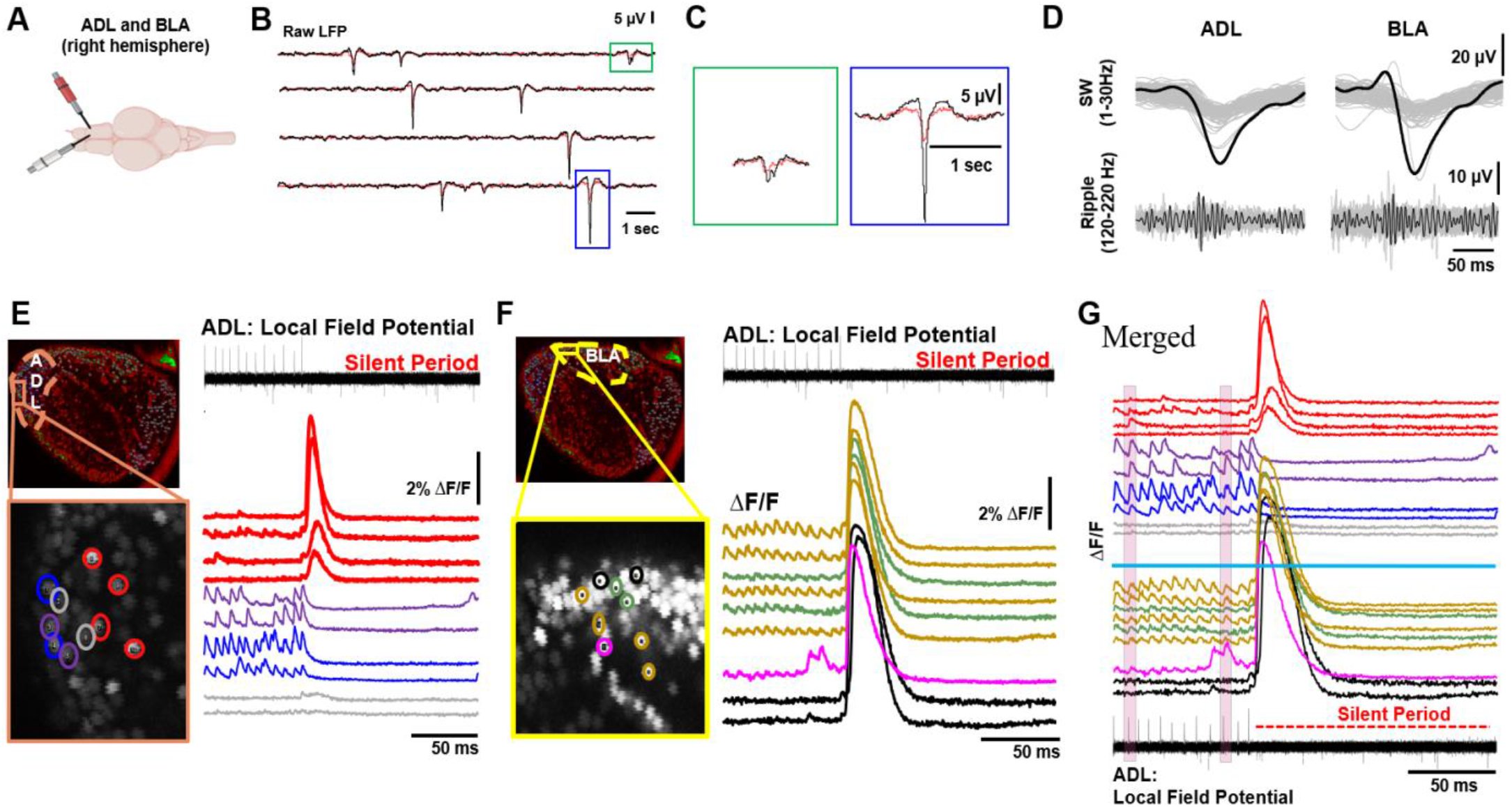
BLA exhibits LFP activity that is coupled to ADL SWR activity. Single cell neuronal activity contributing to SW events is variable. (a) Representative image of electrode placement in both the ADL (red) and BLA (white). (b) Representative superimposed LFP traces (1-1000 Hz) between ADL (red trace) and BLA (blue trace). (c) Zoomed in individual LFP oscillation from (b). (D=d) Representative SWR complex from the ADL and the BLA. (e & f) Representative image of the telencephalon of a *Tg(elevI3:Hsa.H2B:GCaMP6s*) juvenile zebrafish with the ADL (e) or BLA (f) labeled and zoomed in to show ROIs for single cells, along with alignment of single cell activity (color matched) to LFP from the ADL. (g) Merged single cell activity, separating the ADL (top) from the BLA (bottom) by the solid blue line, aligned to ADL LFP. Examples of coupling between neuronal firing in these areas and LFP activity are shown in the pink shaded areas.

**Figure 5.**
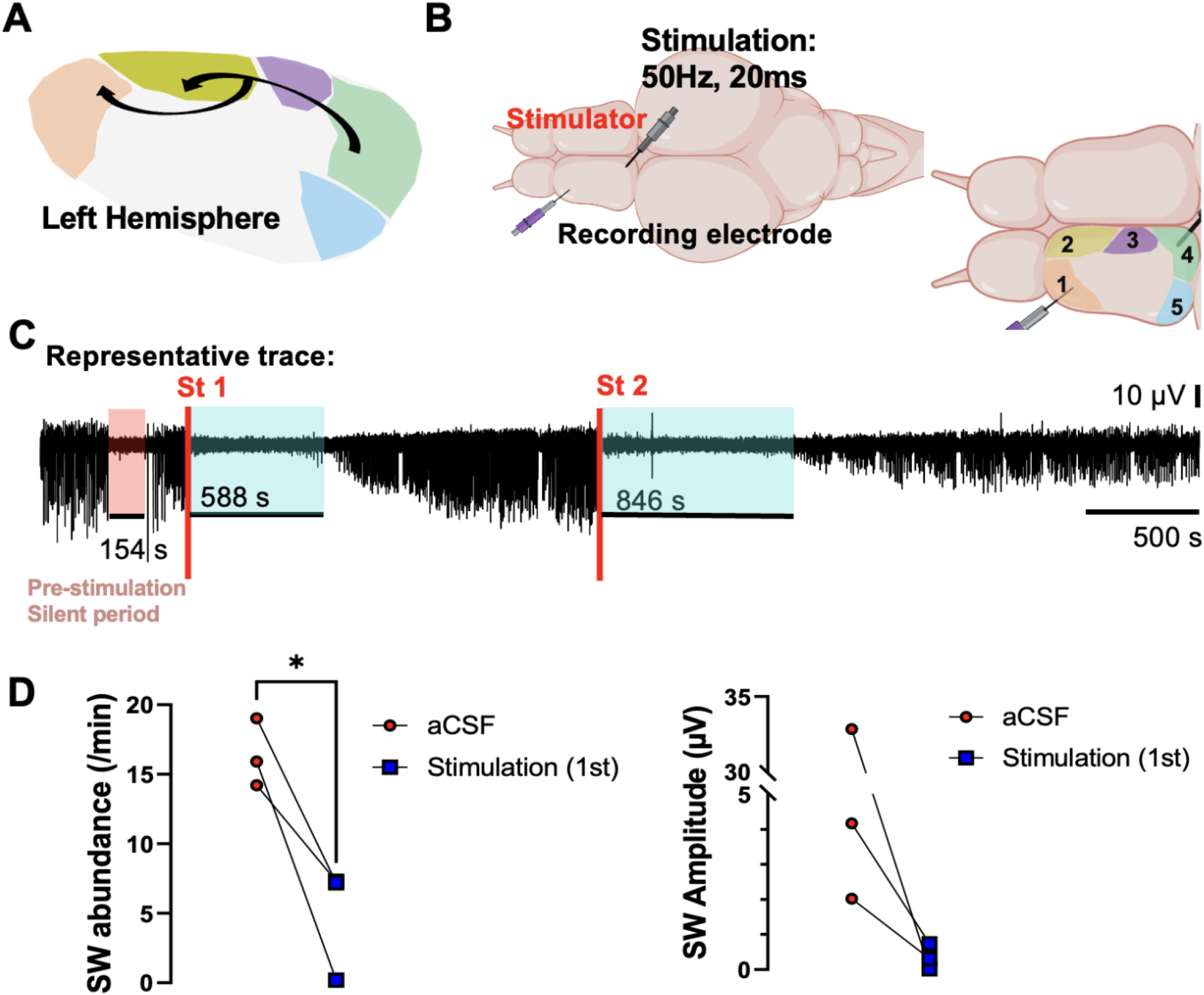
Caudal telencephalic stimulation of putative cholinergic neurons suppresses ADL SW events. (a) Proposed model for the generation of silent periods with activation of region 4 – putative cholinergic neurons. (b) Representative brain showing the placement of the stimulating electrode (region 4) and recording electrode (ADL – region 1). (c) Representative LFP trace showing the start of the stimulation and the resulting suppression of LFP activity (shaded blue area). After suppression the activity comes back and is further suppressed by a second stimulation. Of note: the trace had a silent period before stimulation. (d) Shows the quantification of both SW abundance and the SW amplitude both of which were decreased within the first 10 minutes after the first stimulation.

**Figure 6.**
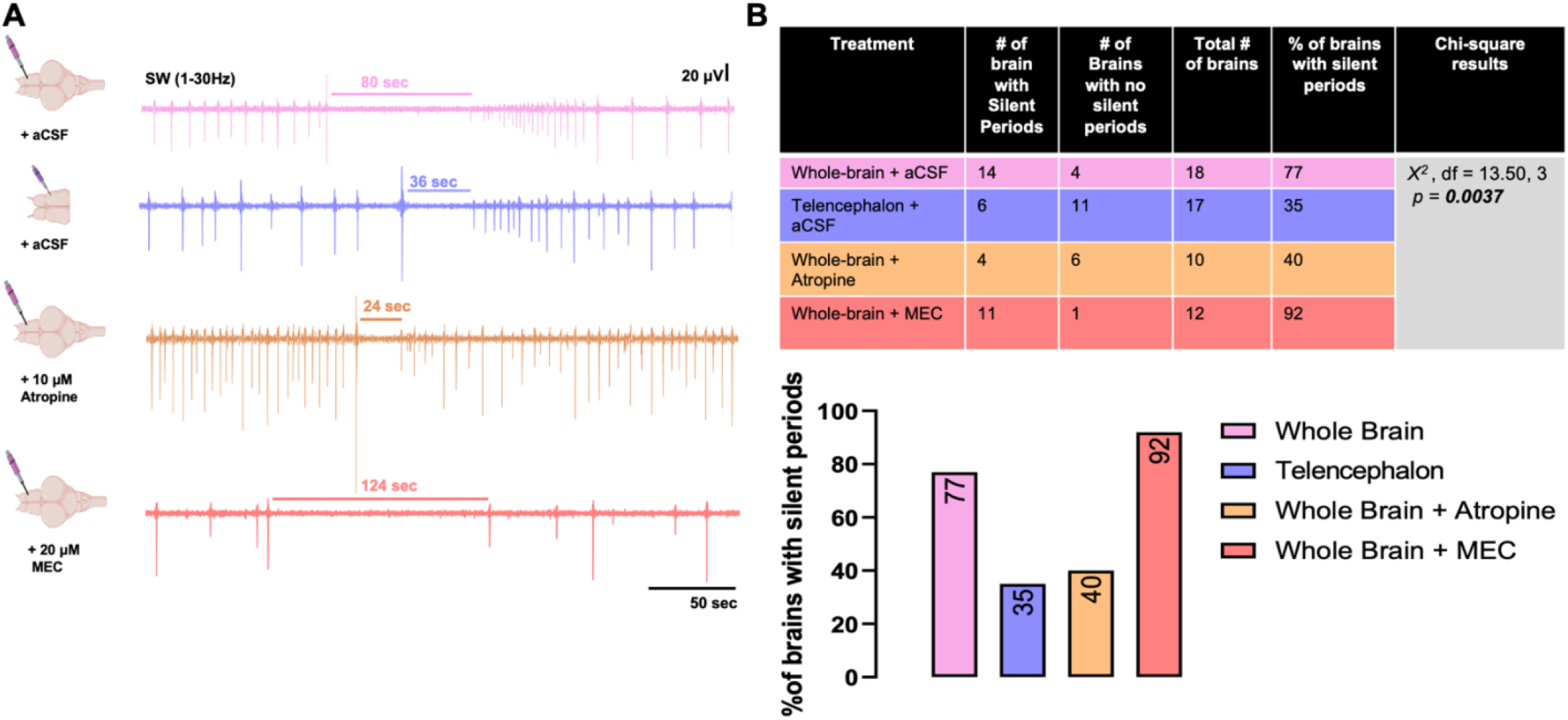
Silent periods are modulated by muscarinic receptors and can be generated intrinsically in the telencephalon. (a) Representative filtered SW events containing silent periods for whole-brain in aCSF, telencephalon only in aCSF, whole-brain + 10 μM atropine, and whole-brain + 20 μM MEC. (b) Quantification of the number of brains containing at least one silent period per recording and bar graph showing the corresponding percent change. Chi-square *X*^2^) = 13.50, df = 3, *p* = 0.0037.

**Figure 7.**
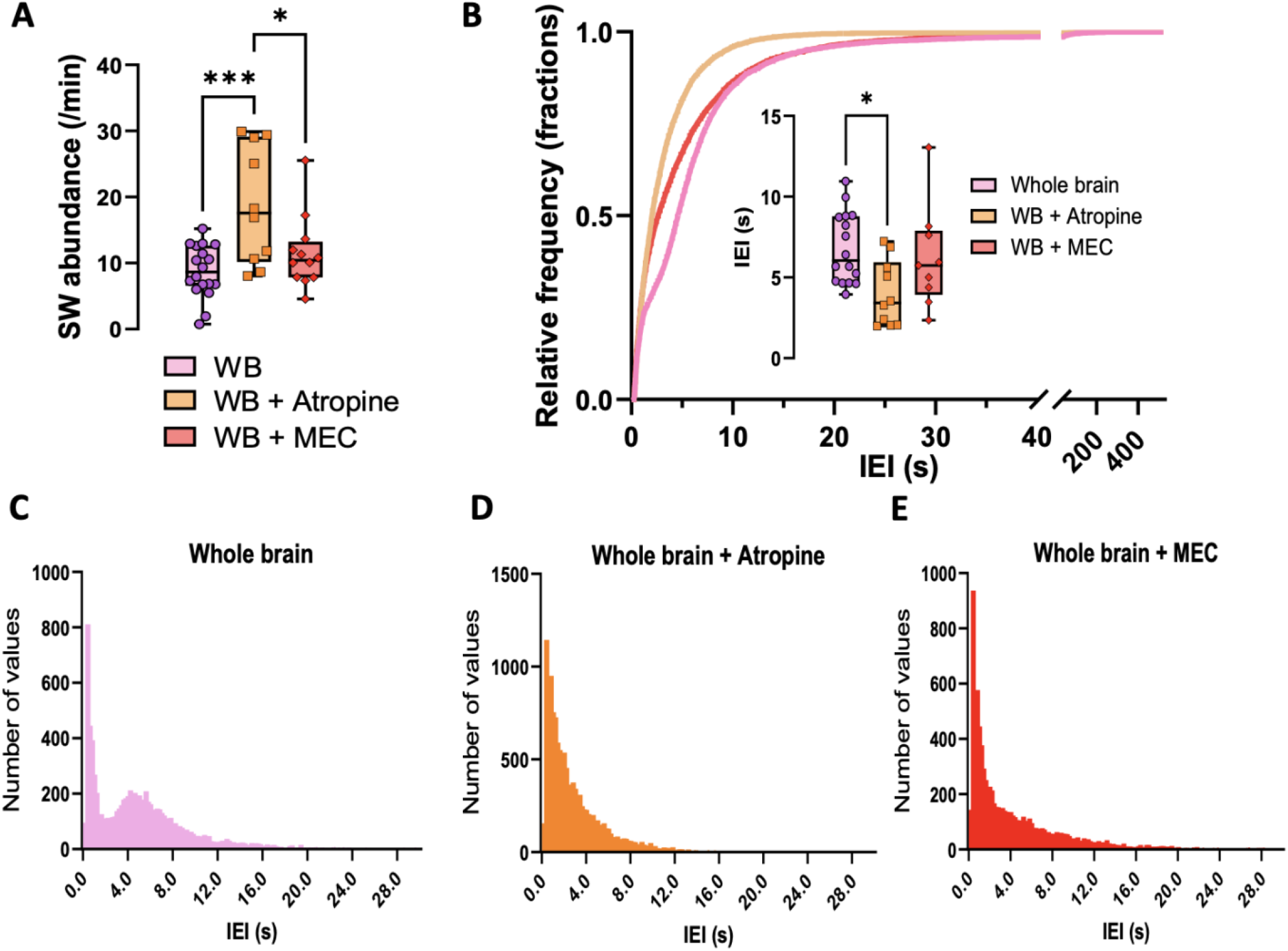
SW events are modulated by the cholinergic system. (a) 30 minutes exposure to 10 μM atropine significantly increased the number of SW events compared to whole-brains in aCSF, whereas exposure to 20 μM MEC did not. (b) Both atropine and MEC shifted the IEI cumulative frequency to the left with atropine showing a significant decrease in the average IEI compared to whole-brain in aCSF. (c-e) Histogram frequencies of IEI with 0.2 bin size and same n as in (b).

### Biorender license

Cartoon images of the zebrafish brain were made with Biorender.com with *Agreement Number: PU24YGDOA2*.

### Code Accessibility

All code is open-source and available in public repositories, including versions under development (Github) as well as archival copies used for this manuscript (Zenodo). MATLAB R2022a functions are available at https://github.com/acaccavano/SWR-Analysis (archival copy: DOI: 10.5281/zenodo.7490166). ImageJ (FIJI) macros are available at https://github.com/acaccavano/deltaFoF (archival copy: DOI: 10.5281/zenodo.362513).

## RESULTS

### SWR events in juvenile zebrafish are locally generated and maintained in the telencephalon

We aimed to characterize the spatial generation, maintenance, and modulation of SW/SWR events in the ADL and BLA in the juvenile zebrafish which also enabled us to record calcium transients at single cell resolution in the entire telencephalon with a confocal microscope using a 20x lens. We previously showed that LFP recordings from the ADL region of the telencephalon of juvenile zebrafish display SW events with above background signal-to-noise ratio after 30 minutes of recovery in oxygenated aCSF at room temperature (Blanco & Conant, 2020). Building upon this observation, we applied dual LFP recordings from the ADL of both hemispheres in a whole-brain preparation and showed that both the left and right hemispheres have spontaneous time-locked LFP signal oscillations (Figure 1A-C). Given that in the mammalian hippocampus SWRs are locally generated and sustained in the absence of intact extra-hippocampal afferent connections (Buzsáki, 1986; Maier et al., 2003), we sought to better understand the spatial generation and maintenance of SWR events in the juvenile zebrafish brain. Therefore, we recorded LFP from whole-brains and de-tectomized (telencephalon only) preparations (Figure 1D). Like whole-brain recordings, de-tectomized brains exhibited SWR events composed of a SW (1-30 Hz) with an occasional embedded fast oscillatory ripple (120-220 Hz) (Figure 1E), suggesting their local generation within the zebrafish telencephalon. Quantification of these events (Figure 1F) reveals a significant increase in the rate of events per minute in de-tectomized preparations (mean = ~14) compared to whole-brain (mean = ~11) (Unpaired Student’s t-test; t_(33)_ = 2.279, p = 0.0293; whole-brain n = 18; de-tectomized n = 17). A trend towards a decrease in the inter-event-interval (IEI) in de-tectomized brains was observed though it did not reach significance (Unpaired Student’s t-test; t_(31)_ = 1.693, p = 0.1004; whole-brain n = 16; de-tectomized n = 17). The amplitude of these events was not changed (Unpaired Student’s t-test; t_(28)_ = 0.9252, p = 0.3628; whole-brain n = 16; de-tectomized n = 14); however, removing the tectum and other extra-telencephalic areas significantly increased their duration (Unpaired Student’s t-test; t_(33)_ = 3.507, p = 0.0013; whole-brain n = 18; de-tectomized n = 17). In our recordings we observed a decrease in the percent of SWR events compared to SW events given that some SW did not exhibit a ripple event (Figure 1G – left panel). Additionally, some SW events had an embedded doublet ripple (Figure 1G – right panel), previously shown to be associated with dendritic summation in rodents (Judák et al., 2022). As a result, we based further analysis on our quantitatively and qualitatively analysis on SW events alone. Together, the data show that SWR events are intrinsically generated and maintained within the telencephalon and that some of their properties are also regulated by extra-telencephalic afferents.

### SWs in juvenile zebrafish are modulated by AMPA and GABA_A_ but not NMDA signaling

To assess whether ADL-specific SW events are modulated by glutamate and GABA neurotransmission as previously described in mammals and whole-brain adult zebrafish (Behrens et al., 2005; Maier et al., 2003; Schlingloff et al., 2014; Vargas et al., 2012; Wu et al., 2005), we perfused whole-brain preparations with 30 μM AP-5, an NMDA receptor antagonist; 25 μM AP-5 + 5 μM NBXQ, an AMPA and kainate receptor antagonist for glutamatergic signaling elimination; and 30 μM bicuculline, a GABA_A_ receptor antagonist. Constant bath perfusion (30 minutes) with 30 μM AP-5 did not significantly affect the average abundance of SW events (paired t-test, t_(7)_ = 0.2984, p = 0.7741, n = 8) (Figure 2A-C). Neither the amplitude nor the duration were significantly changed (paired t-tests, t_(7)_ = 0.4016, p = 0.7000; t_(7)_ = 0.8628, p = 0.4168, respectively); however, the addition of 5 μM NBQX to the bath solution in the presence of 25 μM AP-5 significantly decreased SW events from an average of approximately 10 events per minute to 1 event per minute (paired t-test, t_(3)_ = 4.638, p = 0.0189, n = 4) (Figure 2D-E). Although results were not significant, exposure to NBQX/AP-5 showed a trend towards a decrease in both the amplitude (paired t-test; t_(3)_ = 3.138, p = 0.0518) and the duration (paired t-test; t_(3)_ = 2.391, p = 0.0966) of SW events.

Since locally generated spontaneous SW events are the result of the interplay between excitation and inhibition (Buzsáki, 2015; Eller et al., 2015; Schlingloff et al., 2014; Sipilä et al., 2006; Ylinen et al., 1995), we investigated whether the GABA_A_ receptor antagonist bicuculline (30 μM) affected SWs in whole-brain juvenile zebrafish. After perfusion with bicuculline, there was a significant increase by approximately 50% in the occurrence of spontaneous events (paired t-test; t_(6)_ = 2.643, p = 0.0384, n = 7) (Figure 2G) with a modified intermittent burst activity after filtering in the low-frequency band (1-30 Hz). This is similar to what was previously observed in whole-brain adult zebrafish with a lower dose (10 μM) (Vargas et al., 2012). This modified pattern was composed of bursting events with varied amplitudes that, when averaged across all recording events, showed no significant difference in amplitude between bicuculine-exposed and aCSF-exposed brains (paired t-test, t_(6)_ = 0.2456, p = 0.8142) (Figure 2H inset). Plotting the cumulative distribution for the amplitude (Figure 2H) (aCSF n = 2179, 30 μM bicuculline n = 4052) revealed a significant difference between aCSF and bicuculline-perfused brains (Kolmogorov-Smirnov D test = 0.4713, p < 0.0001). A 2-way ANOVA showed that bicuculline significantly shifted the amplitude of SW events to the left (green arrow) (Drug: F_1, 14_ = 0.08520, p = 0.7746, Bin: F_86, 1204_ = 458.8, p < 0.0001, Drug x Bin: F_86, 1204_ = 1.534, p = 0.0017).

### *Ex vivo* whole-brain preparations exhibit silent period events with no detectable LFP

Sleep-like signatures have been previously shown in the zebrafish. Specifically, periodic slow bursting sleep alternates with propagating wave sleep, which is characterized by an overall reduction in neuronal activity. The latter follows a caudal-to-rostral activation of neurons located in the telencephalic midline in the dorsal pallium of sleep-deprived zebrafish (Leung et al., 2019). In our LFP recordings from the ADL of whole-brain *ex vivo* preparations, we also observed periods with no detectable LFP signal (no SW) ranging from 20 to over 150 seconds in duration, followed by a return of detectable LFP activity (Figure 3A). Most of these events occurred after a high amplitude SW event (purple arrow) with a pattern of subsequent activity that was variable. To understand the origin of silent periods in *ex vivo* whole-brain preparations, we simultaneously recorded LFP events and single cell calcium transient signals using the *Tg(elevI3:Hsa:H2B:GCaMP6s*) zebrafish line. Given the temporally coordinated firing of both telencephalic hemispheres (Figure 1A-C), most of our recordings and analyses were done in the right telencephalon (Figure 3B – top panel). We analyzed the calcium transients resulting from neuronal activity at the regional and single cell levels in two different planes of the zebrafish telencephalon (Figure 3B): the dorsal (D – pink) and the ventral (V – blue) side with a difference in depth of approximately 30-50 μm. At each plane, we selected regions of interest (ROIs) based on the activation pattern during SWs events and silent periods observed under the confocal microscope (numbered and color-coded in Figure 3B). We further selected single cell ROIs within these clusters. We then determined the change in fluorescence over total fluorescence (ΔF/F) for each ROI as previously shown (Caccavano et al., 2020) with some modifications to accommodate the zebrafish data (*see methods*).

Alignment between ADL LFP events and calcium transients from selected ROIs (Figure 3 C1-C2) in the dorsal telencephalon showed spontaneous activity in region 1 (ADL) and region 2 (BLA) that were time-locked during the pre-silent period (Figure 3 C2). Strikingly, using single cell calcium transients, we observed that before a silent period event, the telencephalon underwent a massive activation possibly caused by depolarization (measured by the change in fluorescence – Figure 3 C2: Depolarization) in selected regions that was followed by a total elimination of LFP activity (Figure 3 C1-C2: Silent period). Single cell calcium transients (Figure 3 C3) were measured as ΔF/F within demarked clustered regions and showed that regions previously not contributing to the generation and/or maintenance of SW events, such as region 4 (green) and 5 (blue) – most caudal ones, were involved in the silencing of SW events.

Ventrally, when we examined the pre-chosen clusters (like in the dorsal area), we observed that the total fluorescence activity increased matching that of LFP recordings in ADL and BLA (Figure 3 D1-D3). A similar pattern was shown for the silencing of SW events with neuronal activation going from a caudal to rostral direction (Figure 3 D1-D3). However, we did not see significant changes in calcium transient signals at the single cell level in these ventral areas. This could be the result of the difference in depth of the tissue. The massive activation that culminates in the elimination of SW events in our preparations (dorsal and ventral) typically resulted from a stereotypical pattern of neuronal activity starting at the caudal region of the telencephalon (region 4) and propagating rostrally (Figure 3E). This type of caudal-to-rostral propagation often involved the activation of the telencephalic midline (Supplemental video 2). Though a more detailed analysis including neuronal tracing is needed to better decipher the propagation of caudal-rostral neuronal activity recruitment leading to silent periods, here we propose a working model for the silencing of SW events in the dorsal side of the telencephalon (Figure 3F). We posit that region 4, previously shown through immunohistochemistry to house cholinergic neurons (Clemente et al., 2004), receives input from other areas of the brain which consequently increases the cholinergic tone in the telencephalon and leads to a silencing of SW events in both the ADL and the BLA. We further propose that this caudal region of putative cholinergic neurons may project to both the ADL and BLA via collaterals. Though region #3 is demarcated as an isolated region, it may be part of the amygdaloid complex (Porter & Mueller, 2020) and can explain the single cell activity observed in this region (Figure 3 C3 purple).

### SW event coupling between ALD and BLA in *ex vivo* juvenile zebrafish brains

Previous studies in murine and humans have shown the occurrence of SWR events in the BLA and their coupling to SWR events in the hippocampus (Cox et al., 2020; Paré, 2002; Perumal et al., 2021; Ponomarenko et al., 2003; Popescu & Paré, 2011). Given that the BLA of juvenile zebrafish showed calcium transient neuronal activity that was time-locked to the activity in the ADL (Figure 3 and Supplemental video 1), we next asked if the zebrafish BLA had detectable LFP activity and consequently SW/SWR events. We performed dual LFP recordings from the ADL and BLA (Figure 4A-C) and showed that the zebrafish BLA exhibits LFP activity and that this activity is coupled to ADL LFP activity. We further demonstrated that spontaneously occurring LFP activity in the BLA exhibited events composed of a SW event (1-30 Hz) with a superimposed ripple oscillation (120-220 Hz) (Figure 4D).

We then aligned single cell calcium transient recordings from the ADL (Figure 4E) and the BLA (Figure 4F), to LFP recorded specifically from the ADL to further show time-locked activity between single cells in both regions to ADL LFP recordings (Figure 4E-F). The alignment revealed that not all neurons in the ADL or BLA contribute to a SW event as measured by LFP in the ADL. This is a key similarity between murine hippocampal slices in which only up to 50% of pyramidal neurons contribute to a single SWR event (Ellender et al., 2010; Evangelista et al., 2020; Hajos et al., 2013, p.; Ylinen et al., 1995). Merging of single cell recordings from both brain regions (Figure 4G) corroborates the coupling between neuronal firing in these areas and the generation of a SW event (pink shaded areas in G are examples of coupling). Though not all SW events have an embedded ripple, the high temporal coordination in the LFP activity between the ADL and BLA will result in the coupling of SWR events in these two brain regions similar to that in mammals.

### Stimulation of region 4, containing putative cholinergic neurons, suppresses SW events

Our data suggested (Figures 3 and 4) that silent periods preceded a massive activation of neurons in the telencephalon starting in region #4, which contains putative acetylcholinesterase positive neurons (Clemente et al., 2004). This region is known as the posteromedial pallial amygdala (PMPa) (Porter & Mueller, 2020), and is functionally connected to the BLA (Northcutt, 2008; von Trotha et al., 2014), perhaps explaining some single cell activation in this area during SW events. To corroborate the involvement of the PMPa in silencing SW events in the ADL, we stimulated this region and recorded LFP activity in the ADL (Figure 5A-B). A single stimulation suppressed SW event abundance (paired t-tests, t_(2)_ = 4.507, p = 0.0459, n = 3) and decreased their amplitude (t_(2)_ = 1.254, p = 0.3366, n = 3) despite the later not reaching significance (Figure 5D). In the ADL, the suppression time of SW events was variable (blue shaded areas) with returning events (Figure 5C). Interestingly, we observed that a second stimulation after LFP activity returned to baseline, increased the duration of the suppression, and decreased the amplitude of events compared to the first stimulation. We furthered observed that stimulation of PMPa induced suppression of LFP that was longer than endogenous silent period events (red shaded area).

### Muscarinic receptor antagonism decreases silent periods

We investigated the involvement of the cholinergic system in modulating silent periods and SW events by recording LFP from whole-brains treated with atropine (10 μM), and Mecamylamine (MEC – 20 μM). In the zebrafish, most cholinergic nuclei are located in the diencephalon, mesencephalon and rhombencephalon (Clemente et al., 2004; Toledo-Ibarra et al., 2013). We then reasoned that whole-brain preparations with an intact cholinergic system would exhibit more silent periods than recordings from de-tectomized preparations with a compromised cholinergic system. Silent periods were observed in all recordings (whole-brain in aCSF (n = 18), de-tectomized preparations (n = 17), whole-brains + atropine (n = 10), and whole-brains + MEC (n = 12)) (Figure 6A); however, compared to whole-brain (77% of brains had at least one silent period), de-tectomized preparations had a 2X reduction in the number of recordings exhibiting at least one silent period (Figure 6B). Atropine significantly decreased the number of silent periods within a whole-brain preparation when compared to controls (40% reduction) but was not significantly different from de-tectomized brain preparations. However, MEC did not decrease the number of brains exhibiting a silent period compared to whole-brain recordings. Together, the data show that muscarinic receptors and extra-telencephalic afferents are involved in the modulation of silent periods, and additionally, these findings suggest that silent periods are locally regulated.

### Muscarinic receptor antagonist and agonists differentially modulate SW events

Rodent studies show that blocking muscarinic receptors significantly increases SWR events while enhancing muscarinic activity abolishes SWR events (Y. Zhang et al., 2021). These findings coupled with our result that atropine, a muscarinic antagonist, decreases the number of brains with at least one silent period led us to hypothesize that the introduction of atropine would increase the number of SW events compared to whole-brains. Indeed, we found that atropine increased the number of SW events per minute (aCSF mean = 8.79) approximately two-fold when compared to whole-brains in aCSF (atropine mean = 18.78) (F_(2,37)_ = 9.123, p = 0.0006 one-way ANOVA with Tukey’s multiple comparisons test: p = 0.0004) (Figure 7A). Using the non-competitive nicotinic antagonist MEC, however, did not significantly change the abundance of SW events (mean = 11.96) when compared to whole-brains in aCSF (p = 0.4435). A shift to the left was observed in the IEI cumulative frequencies from whole-brains treated with both atropine and MEC (single events were pulled from recordings in (Figure 7A), whole-brain in aCSF n = 9293, whole-brain + atropine n = 13 400, whole-brain + MEC n = 9008) with a significant average decrease only in brains treated with atropine (Figure 7B – inset) (F_(2,32)_ = 4.121, p = 0.0256 one-way ANOVA with Tukey’s multiple comparisons test) (p = 0.0213). Interestingly, individual IEI histograms revealed an IEI bi-modal distribution that was abolished with both atropine and MEC (Figure 7C-E).

## DISCUSSION

In the present study we show that SWR events recorded from the ADL are intrinsic to the telencephalon, though their abundance and duration can be controlled by extra-telencephalic afferents. Additionally, SWR events occur in the BLA and concomitant single cell calcium imaging and LFP recordings demonstrated that SW events in the ADL are coupled to BLA SW events. Moreover, calcium imaging of whole-brains demonstrated that SWs in both the ADL and BLA are endogenously and spontaneously silenced (silent periods) by the activation of a more caudal population of putative cholinergic cells, region 4 – PMPa. Electrical activation of this region silenced SW events in the ADL. Exposure to atropine decreased the number of brains with at least one silent period and increased the abundance of SW events in the ADL. These data contribute to our understanding of neuronal population dynamics in the zebrafish brain and highlight its advantage for the simultaneous recording of SWs and single cell activity in different brain regions important for learning and memory consolidation. The data also suggest a remarkable conservation of regional SW coupling and cholinergic modulation from mammals to teleost.

### Intrinsically generated SWRs in the ADL

Previously, (Vargas et al., 2012) established the occurrence of SWR events in the ADL – hippocampus homologue – of whole-brain adult zebrafish showing that they had similar features to mammalian SWR events. In the present study we show these events in whole-brain juvenile zebrafish, similar to that of rodents and adult zebrafish (Behrens et al., 2005; Maier et al., 2003; Schlingloff et al., 2014; Vargas et al., 2012; Wu et al., 2005), are modulated by AMPA and GABA_A_, but not NMDA signaling (Figure 2). We corroborated the induction of intermittent burst of spontaneous activity after perfusion with a high dose of bicuculline (30 μM) previously reported in adult zebrafish and murine preparations (Kim et al., 2004; Ogawa et al., 1991; Vargas et al., 2012). A finding of the present study is the time-locked occurrence of SW events between hemispheres and their intrinsic generation and maintenance in the telencephalon (Figure 1). Interestingly, recordings from sleeping Sprague-Dawley rats have previously shown asynchrony between SWR events from the left and right hippocampus due to the lateralization of spatial memories (Villalobos et al., 2017). Thus, future studies may be warranted to determine whether zebrafish ADL SW events are hemispherically uncoupled in select behavioral states. It is important to note that other zebrafish brain regions such as the habenula do show effects of laterality.

Though zebrafish have approximately seven-fold fewer neurons than rodents, as in murine hippocampal slices, we observed that not all neurons in the SW-generating vicinity participate in the genesis and maintenance of a SW event (Figure 4E). Additionally, the activation of neurons during a single SW event does not seem dependent on their direct proximity to each other as neighbor neurons in the ADL behaved differently during a SW event (Figure 4E – grey versus blue or blue and purple). We can speculate that the differential activation of these neurons may resemble the re-activation of neuronal ensembles where not all cells within an area form part of the encoded memory. It has been previously shown that neighbor neurons that participate in a given ensemble do not need to be in direct contact with one another (Minatohara et al., 2016; Wagatsuma et al., 2018).

SW events recorded from the ADL of de-tectomized preparations showed an increase in their abundance and duration (Figure 1F). One explanation for the increase in SW events in the recordings of de-tectomized preparations is an overall decrease in the number of preparations containing a silent period (Figure 6). This pattern is also observed in whole-brains exposed to atropine (discussed in more detail later), which hints at the modulation of SW events by extra-telencephalic afferents. For instance, it is widely accepted that SWR events in mammals are modulated by different subcortical areas including the septum (Nicoll, 1985) and hypothalamus (Vicente et al., 2020). However, the dialogue between the hippocampus and other cortical areas, including the somatosensory cortex during specific types of memory formation (Tukker et al., 2020), cannot be discounted as a potential modulator of SW events in the zebrafish brain.

Future research should investigate the involvement of the zebrafish habenula in controlling silent periods and the abundance and duration of SWs. In the zebrafish, the habenula is located between the telencephalon and the tectum and can be damaged in de-tectomized brains. In the fish, the habenula is additionally shown to house choline acetyltransferase positive neurons (Cheng et al., 2014) which play a critical role in modulating SW events. In rodents, the activity of the habenula has been shown to be phase-locked to theta oscillations in the hippocampus and its silencing modulates hippocampal-dependent spatial recognition tasks (Görlich et al., 2013; Goutagny et al., 2013). This study suggested functional connectivity between the hippocampus and the habenula, specifically the same circuit that gives rise to SW events. The habenula also exerts inhibitory control over brain areas associated with the dopaminergic system (Lecourtier et al., 2008). Indeed, the dopaminergic system, which adds saliency to memories, has been shown to enhance SW events during immobility in rodents (Ambrose et al., 2016; Fuchsberger & Paulsen, 2022; Loren, 2009).

### SW/SWR events in the BLA and their coupling to ADL SW/SWR events

In mammals, diverse cortical/subcortical regions also exhibit synchronous neuronal oscillations, including sleep spindles during stage 2 sleep or SWR events during slow wave sleep, that are coupled to hippocampal SWR events and important to consolidation of different memory types (Eller et al., 2015; Fell et al., 2001; Fries, 2005; Ji & Wilson, 2007; Perumal et al., 2021; Skelin et al., 2019, 2021; Vaz et al., 2020; Wilber et al., 2017). Specifically, in human studies, the BLA has been shown to exhibit SWR events (Perumal et al., 2021) that are coupled to hippocampal SWRs for the consolidation of hippocampal-dependent memories with emotional valence (Cox et al., 2020). In the present study, we present evidence for the generation of SWR events in the BLA of juvenile zebrafish. As in the ADL, a subset of SW events exhibits a superimposed ripple (120-220 Hz) (Figure 1E) and not all neurons contribute to the SW event (Figure 4F). An additional novel finding is the temporal coupling between ADL and BLA SW events as detected by simultaneous LFP and calcium imaging recordings (Figure 4). This supports the possibility that these two structures work in tandem to facilitate the consolidation of emotional memories in zebrafish as they do in mammals through the coupling of SWR events.

In rodents an enriched environment enhances SWR amplitude in subsequently prepared *ex vivo* hippocampal slices (Landeck et al., 2021), and in humans learning just prior to sleep increases slow wave sleep-associated SWR abundance (Eschenko et al., 2008). Aversive learning associated coupling of hippocampal and amygdala activity in rodents is also observed in subsequent slow wave sleep (Girardeau et al., 2017). Thus, it would be of interest to determine whether an experience with emotional valence, such as novelty or fear learning, would increase the abundance of SWRs occurring simultaneously in the ADL and BLA in future studies. The zebrafish model, which facilitates combined electrophysiological recordings and simultaneous visualization of calcium responses in diverse brain regions, would lend itself well to the study of varied types of memory consolidation during physiological and pathological states.

### Cholinergic modulation of SW events and silent periods

The occurrence of SWR events is modulated by cholinergic tone with high cholinergic tone inhibiting SWR events and exploratory memory encoding through the appearance of TGB activity (Buzsaki et al., 1992; Csicsvari & Dupret, 2014; Fisahn et al., 1998; Fuchsberger & Paulsen, 2022; Hasselmo, 1999; Herweg et al., 2020; Konopacki et al., 1987; Li et al., 2019; Ma et al., 2020; Nyhus & Curran, 2010; O’Keefe, 1993; Osipova et al., 2006; Sederberg et al., 2006; Sullivan et al., 2011; Wilson & McNaughton, 1994; S. Zhang et al., 2000; Y. Zhang et al., 2021). A key difference between recordings from *ex vivo* hippocampal murine slices and *ex vivo* whole-brains from zebrafish is that the former lack septo-hippocampal connections. The lack of cholinergic afferents impairs SW modulation and their silencing in these slices. In fact, to stimulate the transition from SW events to TGB activity the addition of exogenous cholinergic agonists to the bath solution is required (Ballinger et al., 2016; Fisahn et al., 1998; Fischer et al., 2014; Hasselmo, 1999). Though we do not yet fully understand the circuitry modulating SWs in zebrafish, many of the cholinergic nuclei are in the diencephalon, mesencephalon, and rhombencephalon (Clemente et al., 2004; Toledo-Ibarra et al., 2013). The presence of acetylcholinesterase (AChE)-positive neurons has also been shown in the telencephalon and the olfactory bulb, and the presence of choline acetyltransferase immunoreactivity (ChAT-ir) fibers in the caudal region of the telencephalon (Clemente et al., 2004). Additionally, it has previously been shown that the transition from slow bursting sleep to propagating wave sleep in zebrafish, which respectively shares characteristics of slow wave sleep and REM sleep in mammals, is stimulated by cholinergic agents (Leung et al., 2019). This modulation is similar to that in mammals, in which acetylcholine tone is off during SW-rich slow wave sleep and on during REM sleep.

In our study, we observed silent events in *ex vivo* whole-brain preparations of juvenile zebrafish recordings, as defined by the absence of SW events in LFP recordings for at least 20 seconds in duration followed by the reappearance of these events with varied amplitude patterns (Figure 3). Single cell calcium imaging revealed that these silent events result from excitation that starts in region 4, housing putative cholinergic neurons (Clemente et al., 2004), and propagates through the entire telencephalon in a caudal-to-rostral fashion. Electrical stimulation of region 4 corroborates the suppression of SW events (Figure 5). Strikingly, previous literature showed that these events, or very similar events called propagating waves, are a signature of REM-like sleep in zebrafish (Leung et al., 2019), which suggests that the *ex vivo* preparation, in an ‘offline’ state, may have entrained sleep state-inducing mechanisms that are independent from external sensory cues. Furthermore, electrical stimulation of this region suppressed SW events for a period that was longer than endogenously generated silent periods. Post electrical stimulation events returned with increasing amplitude on all occasions, a pattern that was observed after endogenous silent periods (Figure 3A – top trace).

To better understand whether silent periods are the result of extra-telencephalic afferents, we recorded SW events from de-tectomized preparations (Figure 6A). Given that SW events are modulated by the cholinergic system and that the zebrafish brain has most of its cholinergic nuclei outside of the telencephalon, we reasoned that the number of de-tectomized brains exhibiting SW events would decrease. Only 50% of de-tectomized brains exhibited at least one silent period, similar to whole-brains treated with atropine (Figure 6 B). MEC, a nicotinic receptor antagonist, however, did not change the number of recordings containing silent periods (Figure 6B) nor did it affect the number of SW events as compared to that in whole-brains in aCSF (Figure 7A). The lack of effect of MEC can be explained by differences in the contribution of nicotinic receptors to the generation and maintenance of SW events in the zebrafish and different routes of exposure (bath exposure like in this study or local exposure) and the kinetics of the receptor and its desensitization. In fact, drugs targeting nicotinic receptors, such as nicotine, have varied effects on the aforementioned parameters (Fischer et al., 2014). Nonetheless, this data implies a conservation in mechanisms for the modulation of SW events by the cholinergic system. Together, these findings support the possibility that *ex vivo* whole-brain preparations may exhibit two different types of silent period events – one which is cholinergic-dependent and is most likely coming from the tectum, and a second which may or may not be cholinergic-dependent and originates within the telencephalon.

In conclusion, the present study shows that SWRs are intrinsic to the telencephalon of zebrafish and that they are modulated by extra-telencephalic afferents. SWs from the right ADL are time-locked to events in the left ADL. The ADL exhibits silent periods with no detectable LFP activity that occur spontaneously following activation of region 4 (PMPa), which contains putative cholinergic neurons. Electrical stimulation of this area corroborated its involvement in silencing SW events. The BLA of zebrafish also exhibits SWR events that are coupled to SWs in the ADL, suggesting hippocampal to cortical/subcortical coupling that may contribute to memory consolidation. Finally, as in rodents, SW events in zebrafish are modulated by the cholinergic system. These data highlight zebrafish, in which whole-brain activity can be easily imaged, as an ideal model to study the coupling of SW events in diverse brain regions using both LFP and single cell calcium transients. Future studies are encouraged to examine SWRs and the modulation of their coupling between diverse brain regions in studies of learning and memory mechanisms, as well as zebrafish models of neurological and psychiatric disorders.

## REFERENCES

Ambrose, R. E., Pfeiffer, B. E., & Foster, D. J. (2016). Reverse Replay of Hippocampal Place Cells Is Uniquely Modulated by Changing Reward. Neuron, 91(5), 1124–1136. https://doi.org/10.1016/j.neuron.2016.07.047

Axmacher, N., Mormann, F., Fernández, G., Elger, C. E., & Fell, J. (2006). Memory formation by neuronal synchronization. Brain Research Reviews, 52(1), 170–182. https://doi.org/10.1016/j.brainresrev.2006.01.007

Ballinger, E. C., Ananth, M., Talmage, D. A., & Role, L. W. (2016). Basal Forebrain Cholinergic Circuits and Signaling in Cognition and Cognitive Decline. Neuron, 91(6), 1199–1218. https://doi.org/10.1016/j.neuron.2016.09.006

Bartoszek, E. M., Ostenrath, A. M., Jetti, S. K., Serneels, B., Mutlu, A. K., Chau, K. T. P., & Yaksi, E. (2021). Ongoing habenular activity is driven by forebrain networks and modulated by olfactory stimuli. Current Biology, 31(17), 3861–3874.e3. https://doi.org/10.1016/j.cub.2021.08.021

Behrens, C. J., van den Boom, L. P., de Hoz, L., Friedman, A., & Heinemann, U. (2005). Induction of sharp wave–ripple complexes in vitro and reorganization of hippocampal networks. Nature Neuroscience, 8(11), 1560–1567. https://doi.org/10.1038/nn1571

Blanco, I., & Conant, K. (2020). Extracellular matrix remodeling with stress and depression: Studies in human, rodent and zebrafish models. European Journal of Neuroscience, ejn.14910. https://doi.org/10.1111/ejn.14910

Bocchio, M., Nabavi, S., & Capogna, M. (2017). Synaptic Plasticity, Engrams, and Network Oscillations in Amygdala Circuits for Storage and Retrieval of Emotional Memories. Neuron, 94(4), 731–743. https://doi.org/10.1016/j.neuron.2017.03.022

Brenet, A., Hassan-Abdi, R., Somkhit, J., Yanicostas, C., & Soussi-Yanicostas, N. (2019). Defective Excitatory/Inhibitory Synaptic Balance and Increased Neuron Apoptosis in a Zebrafish Model of Dravet Syndrome. Cells, 8(10), 1199. https://doi.org/10.3390/cells8101199

Buzsáki, G. (1986). Hippocampal sharp waves: Their origin and significance. Brain Researchs, 398(2), 242–252. https://doi.org/10.1016/0006-8993(86)91483-6

Buzsáki, G. (1996). The Hippocampo-Neocortical Dialogue. Cerebral Cortex, 6(2), 81–92. https://doi.org/10.1093/cercor/6.2.81

Buzsáki, G. (2015). Hippocampal sharp wave-ripple: A cognitive biomarker for episodic memory and planning. Hippocampus, 25(10), 1073–1188. https://doi.org/10.1002/hipo.22488

Buzsaki, G., Horvath, Z., Urioste, R., Hetke, J., & Wise, K. (1992). High-frequency network oscillation in the hippocampus. Science, 256(5059), 1025–1027. https://doi.org/10.1126/science.1589772

Buzsáki, G., Lai-Wo S., L., & Vanderwolf, C. H. (1983). Cellular bases of hippocampal EEG in the behaving rat. Brain Research Reviews, 6(2), 139–171. https://doi.org/10.1016/0165-0173(83)90037-1

Caccavano, A., Bozzelli, P. L., Forcelli, P. A., Pak, D. T. S., Wu, J.-Y., Conant, K., & Vicini, S. (2020). Inhibitory Parvalbumin Basket Cell Activity is Selectively Reduced during Hippocampal Sharp Wave Ripples in a Mouse Model of Familial Alzheimer’s Disease. The Journal of Neuroscience, 40(26), 5116–5136. https://doi.org/10.1523/JNEUROSCI.0425-20.2020

Cheng, R.-K., Jesuthasan, S. J., & Penney, T. B. (2014). Zebrafish forebrain and temporal conditioning. Philosophical Transactions of the Royal Society B: Biological Sciences, 369(1637), 20120462. https://doi.org/10.1098/rstb.2012.0462

Clemente, D., Porteros, Á., Weruaga, E., Alonso, J. R., Arenzana, F. J., Aijón, J., & Arévalo, R. (2004). Cholinergic elements in the zebrafish central nervous system: Histochemical and immunohistochemical analysis: Zebrafish Cholinergic System. Journal of Comparative Neurology, 474(1), 75–107. https://doi.org/10.1002/cne.20111

Colgin, L. L. (2016). Rhythms of the hippocampal network. Nature Reviews Neuroscience, 17(4), 239–249. https://doi.org/10.1038/nrn.2016.21

Cox, R., Rüber, T., Staresina, B. P., & Fell, J. (2020). Sharp Wave-Ripples in Human Amygdala and Their Coordination with Hippocampus during NREM Sleep. Cerebral Cortex Communications, 1(1), tgaa051. https://doi.org/10.1093/texcom/tgaa051

Csicsvari, J., & Dupret, D. (2014). Sharp wave/ripple network oscillations and learning-associated hippocampal maps. Philosophical Transactions of the Royal Society B: Biological Sciences, 369(1635), 20120528. https://doi.org/10.1098/rstb.2012.0528

Ego-Stengel, V., & Wilson, M. A. (2009). Disruption of ripple-associated hippocampal activity during rest impairs spatial learning in the rat. Hippocampus, NA-NA. https://doi.org/10.1002/hipo.20707

Ellender, T. J., Nissen, W., Colgin, L. L., Mann, E. O., & Paulsen, O. (2010). Priming of Hippocampal Population Bursts by Individual Perisomatic-Targeting Interneurons. Journal of Neuroscience, 3θ(17), 5979–5991. https://doi.org/10.1523/JNEUROSCI.3962-09.2010

Eller, J., Zarnadze, S., Bäuerle, P., Dugladze, T., & Gloveli, T. (2015). Cell Type-Specific Separation of Subicular Principal Neurons during Network Activities. PLOS ONE, 10(4), e0123636. https://doi.org/10.1371/journal.pone.0123636

Eschenko, O., Ramadan, W., Molle, M., Born, J., & Sara, S. J. (2008). Sustained increase in hippocampal sharp-wave ripple activity during slow-wave sleep after learning. Learning & Memory, 15(4), 222–228. https://doi.org/10.1101/lm.726008

Evangelista, R., Cano, G., Cooper, C., Schmitz, D., Maier, N., & Kempter, R. (2020). Generation of Sharp Wave-Ripple Events by Disinhibition. The Journal of Neuroscience, 40(41), 7811–7836. https://doi.org/10.1523/JNEUROSCI.2174-19.2020

Fell, J., Klaver, P., Lehnertz, K., Grunwald, T., Schaller, C., Elger, C. E., & Fernández, G. (2001). Human memory formation is accompanied by rhinal–hippocampal coupling and decoupling. Nature Neuroscience, 4(12), 1259–1264. https://doi.org/10.1038/nn759

Feng, P., Becker, B., Zheng, Y., & Feng, T. (2018). Sleep deprivation affects fear memory consolidation: Bi-stable amygdala connectivity with insula and ventromedial prefrontal cortex. Social Cognitive and Affective Neuroscience, 13(2), 145–155. https://doi.org/10.1093/scan/nsx148

Fisahn, A., Pike, F. G., Buhl, E. H., & Paulsen, O. (1998). Cholinergic induction of network oscillations at 40 Hz in the hippocampus in vitro. Nature, 394(6689), 186–189. https://doi.org/10.1038/28179

Fischer, V., Both, M., Draguhn, A., & Egorov, A. V. (2014). Choline-mediated modulation of hippocampal sharp wave-ripple complexes in vitro. Journal of Neurochemistry, 129(5), 792–805. https://doi.org/10.1111/jnc.12693

Fries, P. (2005). A mechanism for cognitive dynamics: Neuronal communication through neuronal coherence. Trends in Cognitive Sciences, 9(10), 474–480. https://doi.org/10.1016/j.tics.2005.08.011

Fuchsberger, T., & Paulsen, O. (2022). Modulation of hippocampal plasticity in learning and memory. Current Opinion in Neurobiology, 75, 102558. https://doi.org/10.1016/j.conb.2022.102558

Ganz, J., Kroehne, V., Freudenreich, D., Machate, A., Geffarth, M., Braasch, I., Kaslin, J., & Brand, M. (2015). Subdivisions of the adult zebrafish pallium based on molecular marker analysis. F1000Research, 3, 308. https://doi.org/10.12688/f1000research.5595.2

Girardeau, G., Benchenane, K., Wiener, S. I., Buzsáki, G., & Zugaro, M. B. (2009). Selective suppression of hippocampal ripples impairs spatial memory. Nature Neuroscience, 12(10), 1222–1223. https://doi.org/10.1038/nn.2384

Girardeau, G., Cei, A., & Zugaro, M. (2014). Learning-Induced Plasticity Regulates Hippocampal Sharp Wave-Ripple Drive. Journal of Neuroscience, 34(15), 5176–5183. https://doi.org/10.1523/JNEUROSCI.4288-13.2014

Girardeau, G., Inema, I., & Buzsáki, G. (2017). Reactivations of emotional memory in the hippocampus–amygdala system during sleep. Nature Neuroscience, 20(11), 1634–1642. https://doi.org/10.1038/nn.4637

Görlich, A., Antolin-Fontes, B., Ables, J. L., Frahm, S., Ślimak, M. A., Dougherty, J. D., & Ibañez-Tallon, I. (2013). Reexposure to nicotine during withdrawal increases the pacemaking activity of cholinergic habenular neurons. Proceedings of the National Academy of Sciences, 110(42), 17077–17082. https://doi.org/10.1073/pnas.1313103110

Goutagny, R., Loureiro, M., Jackson, J., Chaumont, J., Williams, S., Isope, P., Kelche, C., Cassel, J.-C., & Lecourtier, L. (2013). Interactions between the Lateral Habenula and the Hippocampus: Implication for Spatial Memory Processes. Neuropsychopharmacology, 38(12), 2418–2426. https://doi.org/10.1038/npp.2013.142

Hajos, N., Karlocai, M. R., Nemeth, B., Ulbert, I., Monyer, H., Szabo, G., Erdelyi, F., Freund, T. F., & Gulyas, A. I. (2013). Input-Output Features of Anatomically Identified CA3 Neurons during Hippocampal Sharp Wave/Ripple Oscillation In Vitro. Journal of Neuroscience, 33(28), 11677–11691. https://doi.org/10.1523/JNEUROSCI.5729-12.2013

Hashimoto, A., Sawada, T., & Natsume, K. (2017). The change of picrotoxin-induced epileptiform discharges to the beta oscillation by carbachol in rat hippocampal slices. Biophysics and Physicobiology, 14(0), 137–146. https://doi.org/10.2142/biophysico.14.0_137

Hasselmo, null. (1999). Neuromodulation: Acetylcholine and memory consolidation. Trends in Cognitive Sciences, 3(9), 351–359. https://doi.org/10.1016/s1364-6613(99)01365-0

Herweg, N. A., Solomon, E. A., & Kahana, M. J. (2020). Theta Oscillations in Human Memory. Trends in Cognitive Sciences, 24(3), 208–227. https://doi.org/10.1016/j.tics.2019.12.006

Ji, D., & Wilson, M. A. (2007). Coordinated memory replay in the visual cortex and hippocampus during sleep. Nature Neuroscience, 10(1), 100–107. https://doi.org/10.1038/nn1825

Jones, E. A., Gillespie, A. K., Yoon, S. Y., Frank, L. M., & Huang, Y. (2019). Early Hippocampal Sharp-Wave Ripple Deficits Predict Later Learning and Memory Impairments in an Alzheimer’s Disease Mouse Model. Cell Reports, 29(8), 2123–2133.e4. https://doi.org/10.1016/j.celrep.2019.10.056

Joo, H. R., & Frank, L. M. (2018). The hippocampal sharp wave–ripple in memory retrieval for immediate use and consolidation. Nature Reviews Neuroscience, 19(12), 744–757. https://doi.org/10.1038/s41583-018-0077-1

Judák, L., Chiovini, B., Juhász, G., Pálfi, D., Mezriczky, Z., Szadai, Z., Katona, G., Szmola, B., Ócsai, K., Martinecz, B., Mihály, A., Dénes, Á., Kerekes, B., Szepesi, Á., Szalay, G., Ulbert, I., Mucsi, Z., Roska, B., & Rózsa, B. (2022). Sharp-wave ripple doublets induce complex dendritic spikes in parvalbumin interneurons in vivo. Nature Communications, 13(1), 6715. https://doi.org/10.1038/s41467-022-34520-1

Kim, Y.-J., Nam, R.-H., Yoo, Y. M., & Lee, C.-J. (2004). Identification and functional evidence of GABAergic neurons in parts of the brain of adult zebrafish (Danio rerio). Neuroscience Letters, 355(1–2), 29–32. https://doi.org/10.1016/j.neulet.2003.10.024

Konopacki, J., Bruce MacIver, M., Bland, B. H., & Roth, S. H. (1987). Carbachol-induced EEG ‘theta’ activity in hippocampal brain slices. Brain Research, 405(1), 196–198. https://doi.org/10.1016/0006-8993(87)91009-2

Lal, P., & Kawakami, K. (2022). Integrated Behavioral, Genetic and Brain Circuit Visualization Methods to Unravel Functional Anatomy of Zebrafish Amygdala. Frontiers in Neuroanatomy, 16, 837527. https://doi.org/10.3389/fnana.2022.837527

Landeck, L., Kaiser, M. E., Hefter, D., Draguhn, A., & Both, M. (2021). Enriched Environment Modulates Sharp Wave-Ripple (SPW-R) Activity in Hippocampal Slices. Frontiers in Neural Circuits, 15, 758939. https://doi.org/10.3389/fncir.2021.758939

Lecourtier, L., DeFrancesco, A., & Moghaddam, B. (2008). Differential tonic influence of lateral habenula on prefrontal cortex and nucleus accumbens dopamine release. European Journal of Neuroscience, 27(7), 1755–1762. https://doi.org/10.1111/j.1460-9568.2008.06130.x

Leung, L. C., Wang, G. X., Madelaine, R., Skariah, G., Kawakami, K., Deisseroth, K., Urban, A. E., & Mourrain, P. (2019). Neural signatures of sleep in zebrafish. Nature, 571(7764), 198–204. https://doi.org/10.1038/s41586-019-1336-7

Li, P., Geng, X., Jiang, H., Caccavano, A., Vicini, S., & Wu, J. (2019). Measuring Sharp Waves and Oscillatory Population Activity With the Genetically Encoded Calcium Indicator GCaMP6f. Frontiers in Cellular Neuroscience, 13, 274. https://doi.org/10.3389/fncel.2019.00274

Loren, F. (2009). Reward enhances reactivation of experience in the hippocampus. Frontiers in Systems Neuroscience, 3. https://doi.org/10.3389/conf.neuro.06.2009.03.303

Ma, X., Zhang, Y., Wang, L., Li, N., Barkai, E., Zhang, X., Lin, L., & Xu, J. (2020). The Firing of Theta State-Related Septal Cholinergic Neurons Disrupt Hippocampal Ripple Oscillations via Muscarinic Receptors. The Journal of Neuroscience, 40(18), 3591–3603. https://doi.org/10.1523/JNEUROSCI.1568-19.2020

Maier, N., Nimmrich, V., & Draguhn, A. (2003). Cellular and Network Mechanisms Underlying Spontaneous Sharp Wave–Ripple Complexes in Mouse Hippocampal Slices. The Journal of Physiology, 550(3), 873–887. https://doi.org/10.1113/jphysiol.2003.044602

McGaugh, J. L. (2004). THE AMYGDALA MODULATES THE CONSOLIDATION OF MEMORIES OF EMOTIONALLY AROUSING EXPERIENCES. Annual Review of Neuroscience, 27(1), 1–28. https://doi.org/10.1146/annurev.neuro.27.070203.144157

Minatohara, K., Akiyoshi, M., & Okuno, H. (2016). Role of Immediate-Early Genes in Synaptic Plasticity and Neuronal Ensembles Underlying the Memory Trace. Frontiers in Molecular Neuroscience, 8. https://doi.org/10.3389/fnmol.2015.00078

Mölle, M., Yeshenko, O., Marshall, L., Sara, S. J., & Born, J. (2006). Hippocampal Sharp Wave-Ripples Linked to Slow Oscillations in Rat Slow-Wave Sleep. Journal of Neurophysiology, 96(1), 62–70. https://doi.org/10.1152/jn.00014.2006

Nicoll, R. A. (1985). The septo-hippocampal projection: A model cholinergic pathway. Trends in Neurosciences, 8, 533–536. https://doi.org/10.1016/0166-2236(85)90190-0

Norman, Y., Yeagle, E. M., Khuvis, S., Harel, M., Mehta, A. D., & Malach, R. (2019). Hippocampal sharp-wave ripples linked to visual episodic recollection in humans. Science, 365(6454), eaax1030. https://doi.org/10.1126/science.aax1030

Northcutt, R. G. (2008). Forebrain evolution in bony fishes. Brain Research Bulletin, 75(2–4), 191–205. https://doi.org/10.1016/j.brainresbull.2007.10.058

Nyhus, E., & Curran, T. (2010). Functional role of gamma and theta oscillations in episodic memory. Neuroscience & Biobehavioral Reviews, 34(7), 1023–1035. https://doi.org/10.1016/j.neubiorev.2009.12.014

Ogawa, S., Kow, L. M., & Pfaff, D. W. (1991). Effects of GABA and related agents on the electrical activity of hypothalamic ventromedial nucleus neurons in vitro. Experimental Brain Research, 85(1). https://doi.org/10.1007/BF00229989

O’Keefe, J. (1993). Hippocampus, theta, and spatial memory. Current Opinion in Neurobiology, 3(6), 917–924. https://doi.org/10.1016/0959-4388(93)90163-S

O’Neill, P.-K., Gore, F., & Salzman, C. D. (2018). Basolateral amygdala circuitry in positive and negative valence. Current Opinion in Neurobiology, 49, 175–183. https://doi.org/10.1016/j.conb.2018.02.012

Osipova, D., Takashima, A., Oostenveld, R., Fernandez, G., Maris, E., & Jensen, O. (2006). Theta and Gamma Oscillations Predict Encoding and Retrieval of Declarative Memory. Journal of Neuroscience, 26(28), 7523–7531. https://doi.org/10.1523/JNEUROSCI.1948-06.2006

Paré, D. (2002). Amygdala oscillations and the consolidation of emotional memories. Trends in Cognitive Sciences, 6(7), 306–314. https://doi.org/10.1016/S1364-6613(02)01924-1

Perumal, M. B., Latimer, B., Xu, L., Stratton, P., Nair, S., & Sah, P. (2021). Microcircuit mechanisms for the generation of sharp-wave ripples in the basolateral amygdala: A role for chandelier interneurons. Cell Reports, 35(6), 109106. https://doi.org/10.1016/j.celrep.2021.109106

Pignatelli, M., & Beyeler, A. (2019). Valence coding in amygdala circuits. Current Opinion in Behavioral Sciences, 26, 97–106. https://doi.org/10.1016/j.cobeha.2018.10.010

Ponomarenko, A. A., Korotkova, T. M., & Haas, H. L. (2003). High frequency (200 Hz) oscillations and firing patterns in the basolateral amygdala and dorsal endopiriform nucleus of the behaving rat. Behavioural Brain Research, 141(2), 123–129. https://doi.org/10.1016/S0166-4328(02)00327-3

Popescu, A. T., & Paré, D. (2011). Synaptic Interactions Underlying Synchronized Inhibition in the Basal Amygdala: Evidence for Existence of Two Types of Projection Cells. Journal of Neurophysiology, 105(2), 687–696. https://doi.org/10.1152/jn.00732.2010

Porter, B. A., & Mueller, T. (2020). The Zebrafish Amygdaloid Complex – Functional Ground Plan, Molecular Delineation, and Everted Topology. Frontiers in Neuroscience, 14, 608. https://doi.org/10.3389/fnins.2020.00608

Sadowski, J. H. L. P., Jones, M. W., & Mellor, J. R. (2016). Sharp-Wave Ripples Orchestrate the Induction of Synaptic Plasticity during Reactivation of Place Cell Firing Patterns in the Hippocampus. Cell Reports, 14(8), 1916–1929. https://doi.org/10.1016/j.celrep.2016.01.061

Schlingloff, D., Kali, S., Freund, T. F., Hajos, N., & Gulyas, A. I. (2014). Mechanisms of Sharp Wave Initiation and Ripple Generation. Journal of Neuroscience, 34(34), 11385–11398. https://doi.org/10.1523/JNEUROSCI.0867-14.2014

Sederberg, P. B., Schulze-Bonhage, A., Madsen, J. R., Bromfield, E. B., McCarthy, D. C., Brandt, A., Tully, M. S., & Kahana, M. J. (2006). Hippocampal and Neocortical Gamma Oscillations Predict Memory Formation in Humans. Cerebral Cortex, 17(5), 1190–1196. https://doi.org/10.1093/cercor/bhl030

Sipilä, S. T., Schuchmann, S., Voipio, J., Yamada, J., & Kaila, K. (2006). The cation-chloride cotransporter NKCC1 promotes sharp waves in the neonatal rat hippocampus: NKCC1 promotes early sharp waves. The Journal of Physiology, 573(3), 765–773. https://doi.org/10.1113/jphysiol.2006.107086

Skelin, I., Kilianski, S., & McNaughton, B. L. (2019). Hippocampal coupling with cortical and subcortical structures in the context of memory consolidation. Neurobiology of Learning and Memory, 160, 21–31. https://doi.org/10.1016/j.nlm.2018.04.004

Skelin, I., Zhang, H., Zheng, J., Ma, S., Mander, B. A., Kim McManus, O., Vadera, S., Knight, R. T., McNaughton, B. L., & Lin, J. J. (2021). Coupling between slow waves and sharp-wave ripples engages distributed neural activity during sleep in humans. Proceedings of the National Academy of Sciences, 118(21), e2012075118. https://doi.org/10.1073/pnas.2012075118

Stork, O., & Pape, H.-C. (2002). Fear memory and the amygdala: Insights from a molecular perspective. Cell and Tissue Research, 310(3), 271–277. https://doi.org/10.1007/s00441-002-0656-2

Sullivan, D., Csicsvari, J., Mizuseki, K., Montgomery, S., Diba, K., & Buzsaki, G. (2011). Relationships between Hippocampal Sharp Waves, Ripples, and Fast Gamma Oscillation: Influence of Dentate and Entorhinal Cortical Activity. Journal of Neuroscience, 31(23), 8605–8616. https://doi.org/10.1523/JNEUROSCI.0294-11.2011

Sun, Z. Y., Bozzelli, P. L., Caccavano, A., Allen, M., Balmuth, J., Vicini, S., Wu, J.-Y., & Conant, K. (2018). Disruption of perineuronal nets increases the frequency of sharp wave ripple events. Hippocampus, 28(1), 42–52. https://doi.org/10.1002/hipo.22804

Toledo-Ibarra, G. A., Rojas-Mayorquín, A. E., & Girón-Pérez, M. I. (2013). Influence of the Cholinergic System on the Immune Response of Teleost Fishes: Potential Model in Biomedical Research. Clinical and Developmental Immunology, 2013, 1–9. https://doi.org/10.1155/2013/536534

Tukker, J. J., Beed, P., Schmitz, D., Larkum, M. E., & Sachdev, R. N. S. (2020). Up and Down States and Memory Consolidation Across Somatosensory, Entorhinal, and Hippocampal Cortices. Frontiers in Systems Neuroscience, 14, 22. https://doi.org/10.3389/fnsys.2020.00022

Vandecasteele, M., Varga, V., Berényi, A., Papp, E., Barthó, P., Venance, L., Freund, T. F., & Buzsáki, G. (2014). Optogenetic activation of septal cholinergic neurons suppresses sharp wave ripples and enhances theta oscillations in the hippocampus. Proceedings of the National Academy of Sciences, 777(37), 13535–13540. https://doi.org/10.1073/pnas.1411233111

Vargas, R., Jóhannesdóttir, I. b., Sigurgeirsson, B., borsteinsson, H., & Karlsson, K.Æ. (2011). The zebrafish brain in research and teaching: A simple in vivo and in vitro model for the study of spontaneous neural activity. Advances in Physiology Education, 35(2), 188–196. https://doi.org/10.1152/advan.00099.2010

Vargas, R., Þorsteinsson, H., & Karlsson, K. Æ. (2012). Spontaneous neural activity of the anterodorsal lobe and entopeduncular nucleus in adult zebrafish: A putative homologue of hippocampal sharp waves. Behavioural Brain Research, 229(1), 10–20. https://doi.org/10.1016/j.bbr.2011.12.025

Vaz, A. P., Wittig, J. H., Inati, S. K., & Zaghloul, K. A. (2020). Replay of cortical spiking sequences during human memory retrieval. Science, 367(6482), 1131–1134. https://doi.org/10.1126/science.aba0672

Vicente, A. F., Slézia, A., Ghestem, A., Bernard, C., & Quilichini, P. P. (2020). In Vivo Characterization of Neurophysiological Diversity in the Lateral Supramammillary Nucleus during Hippocampal Sharp–wave Ripples of Adult Rats. Neuroscience, 435, 95–111. https://doi.org/10.1016/j.neuroscience.2020.03.034

Villalobos, C., Maldonado, P. E., & Valdés, J. L. (2017). Asynchronous ripple oscillations between left and right hippocampi during slow-wave sleep. PLOS ONE, 12(2), e0171304. https://doi.org/10.1371/journal.pone.0171304

von Trotha, J. W., Vernier, P., & Bally-Cuif, L. (2014). Emotions and motivated behavior converge on an amygdala-like structure in the zebrafish. European Journal of Neuroscience, 40(9), 3302–3315. https://doi.org/10.1111/ejn.12692

Wagatsuma, A., Okuyama, T., Sun, C., Smith, L. M., Abe, K., & Tonegawa, S. (2018). Locus coeruleus input to hippocampal CA3 drives single-trial learning of a novel context. Proceedings of the National Academy of Sciences, 115(2). https://doi.org/10.1073/pnas.1714082115

Wilber, A. A., Skelin, I., Wu, W., & McNaughton, B. L. (2017). Laminar Organization of Encoding and Memory Reactivation in the Parietal Cortex. Neuron, 95(6), 1406–1419.e5. https://doi.org/10.1016/j.neuron.2017.08.033

Wilson, M. A., & McNaughton, B. L. (1994). Reactivation of hippocampal ensemble memories during sleep. Science (New York, N.Y.), 265(5172), 676–679. https://doi.org/10.1126/science.8036517

Wu, C., Asl, M. N., Gillis, J., Skinner, F. K., & Zhang, L. (2005). An In Vitro Model of Hippocampal Sharp Waves: Regional Initiation and Intracellular Correlates. Journal of Neurophysiology, 94(1), 741–753. https://doi.org/10.1152/jn.00086.2005

Ylinen, A., Bragin, A., Nadasdy, Z., Jando, G., Szabo, I., Sik, A., & Buzsaki, G. (1995). Sharp wave-associated high-frequency oscillation (200 Hz) in the intact hippocampus: Network and intracellular mechanisms. The Journal of Neuroscience, 15(1), 30–46. https://doi.org/10.1523/JNEUROSCI.15-01-00030.1995

Zhang, S., José, J. V., & Tiesinga, P. H. E. (2000). Model of carbachol-induced gamma-frequency oscillations in hippocampus. Neurocomputing, 32–33, 617–622. https://doi.org/10.1016/S0925-2312(00)00223-X

Zhang, Y., Cao, L., Varga, V., Jing, M., Karadas, M., Li, Y., & Buzsáki, G. (2021). Cholinergic suppression of hippocampal sharp-wave ripples impairs working memory. Proceedings of the National Academy of Sciences of the United States of America, 118(15), e2016432118. https://doi.org/10.1073/pnas.2016432118

Zhou, H., Neville, K. R., Goldstein, N., Kabu, S., Kausar, N., Ye, R., Nguyen, T. T., Gelwan, N., Hyman, B. T., & Gomperts, S. N. (2019). Cholinergic modulation of hippocampal calcium activity across the sleep-wake cycle. ELife, 8, e39777. https://doi.org/10.7554/eLife.39777

